# Pannexin 1 influences lineage specification of human iPSCs

**DOI:** 10.1101/2021.01.21.427632

**Authors:** Rebecca J. Noort, Grace A. Christopher, Jessica L. Esseltine

**Author notes:** Authors contributed equally (Rebecca J. Noort, Grace A. Christopher). To whom correspondence can be addressed.

## Abstract

Every single cell in the body communicates with nearby cells to locally organize activities with their neighbors and dysfunctional cell-cell communication can be detrimental during cell lineage commitment, tissue patterning and organ development. **Pannexin channels (PANX1, PANX2, PANX3)** facilitate purinergic paracrine signaling through the passage of messenger molecules out of cells. PANX1 is widely expressed throughout the body and has recently been identified in human oocytes as well as 2 and 4-cell stage human embryos. Given its abundance across multiple adult tissues and its expression at the earliest stages of human development, we sought to understand whether PANX1 impacts human induced pluripotent stem cells (iPSCs) or plays a role in cell fate decisions. Western blot, immunofluorescence and flow cytometry reveal that PANX1 is expressed in iPSCs as well as all three germ lineages derived from these cells: ectoderm, endoderm, and mesoderm. PANX1 demonstrates differential glycosylation patterns and subcellular localization across the germ lineages. Using CRISPR-Cas9 gene ablation, we find that loss of PANX1 has no obvious impact on iPSC morphology, survival, or pluripotency gene expression. However, *PANX1* knockout iPSCs exhibit apparent lineage specification bias during 2-dimensional and 3-dimensional spontaneous differentiation into the three germ lineages. Indeed, loss of PANX1 significantly decreases the proportion of ectodermal cells within spontaneously differentiated cultures, while endodermal and mesodermal representation is increased in PANX1 knockout cells. Importantly, *PANX1* knockout iPSCs are fully capable of differentiating toward each specific lineage when exposed to the appropriate external signaling pressures, suggesting that although PANX1 influences germ lineage specification, it is not essential to this process.

**Graphical abstract:** 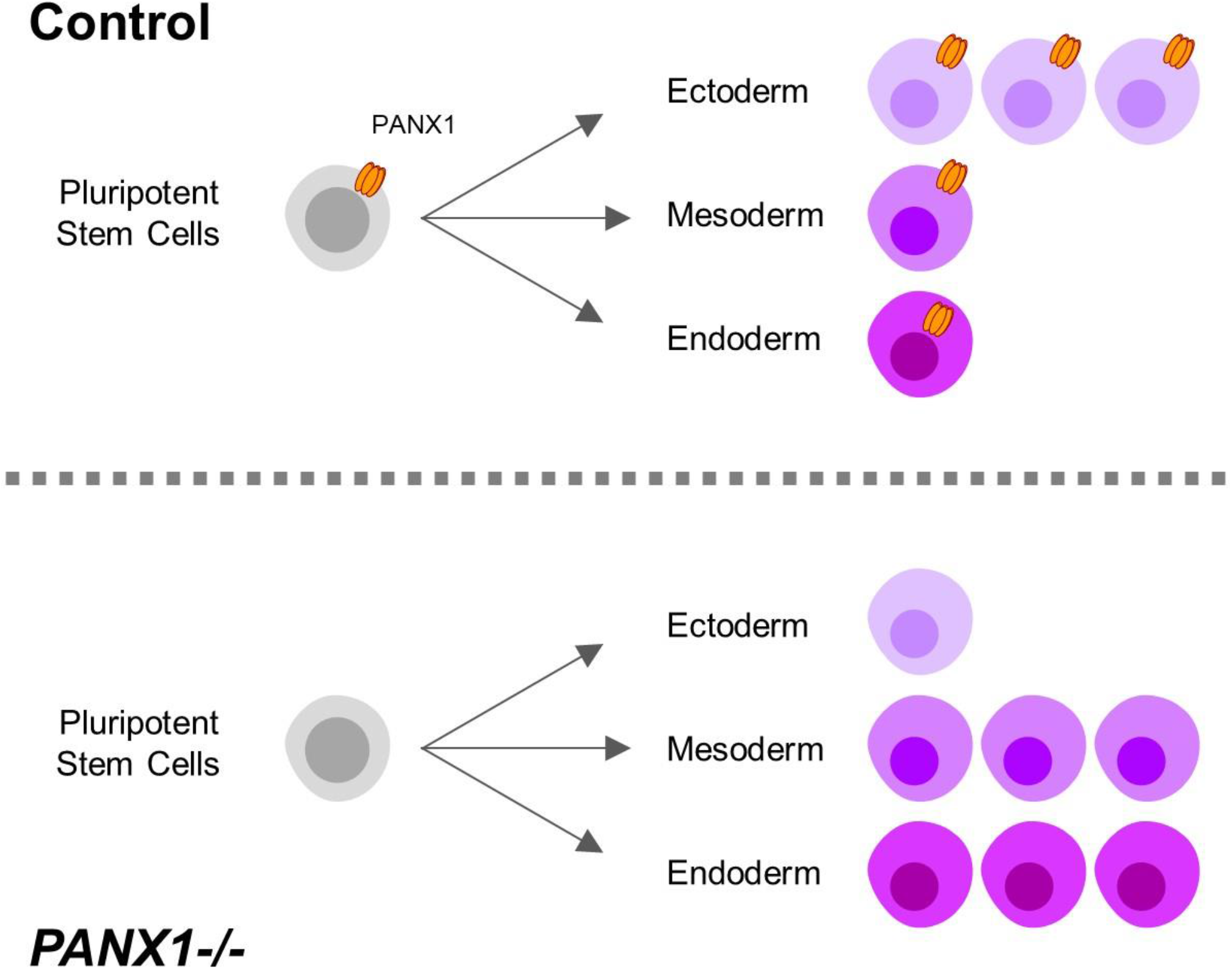

## Introduction

Pluripotent stem cells (PSCs), including embryonic stem cells (ESCs) and induced pluripotent stem cells (iPSCs) are characterized by the ability to self-renew indefinitely and the capacity to differentiate into theoretically any cell type in the embryo (1,2). Differentiation is the process by which stem cells are assigned a cell fate. At each successive stage of lineage commitment cells become more and more specialized and lose the capacity to become other cell types. Terminal differentiation refers to the final stage of cell fate specification, when a cell is locked in form and function. One of the first cell fate decisions in development occurs during gastrulation when the embryo patterns the three embryonic germ layers: ectoderm, endoderm, mesoderm. The ectoderm eventually gives rise to tissues including skin and brain while the endoderm patterns for internal organs and the mesoderm contributes to muscle, bones, and connective tissues. Embryonic germ layer specification can be mimicked *in vitro* through active or passive differentiation paradigms of human PSCs.

Pluripotent stem cells (and epiblast cells in the embryo) maintain their stemness through activation of POU5F1, NANOG, and TGFβ signaling pathways (reviewed in (3)). Gastrulation and the subsequent emergence of the three germ layers from pluripotent stem cells initiates when Activin/Nodal, BMP, and WNT signaling pathways are activated (4). WNT3 signaling enables formation of the primitive streak specification of the bi-potent mesendoderm (3,5,6). The mesoderm subsequently specifies through additional WNT3 pathway activation and BMP signaling, whereas endoderm is specified by elevated activation of Nodal/Activin A signaling pathways (3,7). The ectoderm germ lineage is formed when remaining epiblast cells that did not ingress through the primitive streak are subjected to TGFβ/Nodal and BMP pathway inhibition (3,8). Germ lineage specification can be mimicked *in vitro* through exogenous exposure to the aforementioned signaling molecules in directed differentiation of PSCs. In contrast to directed differentiation, spontaneous differentiation enables passive, cell-guided differentiation and generally results in cell populations from all three germ layers (1,9,10).

Cells do not undertake differentiation in isolation; rather communication among neighboring cells, and between cells and the niche, is essential to ensuring appropriate cell fate specification (11). In addition to secreted factors, small signaling molecules released from cellular channels also play increasingly recognized roles in these processes. For example, purinergic signaling through release of extracellular ATP participates in neural precursor cell and mesenchymal stem cell self-renewal, migration and differentiation (12–15). Pannexin proteins (PANX1, PANX2 and PANX3) form large-pore channels, permeable to ions and metabolites less than 1 kDa in size (16). PANX1 is widely expressed across multiple tissues in the body while PANX2 and PANX3 are more restricted in tissue expression (16). The most well defined role of PANX1 is ATP release (17). In healthy cells PANX1 channels normally remain in a closed state until induced to transiently open in response to a variety of stimuli including membrane deformation, receptor activation, and intracellular calcium release (18–21). PANX1 signaling can also influence the self-renewal and differentiation of multiple somatic (adult) stem cell types including osteoprogenitor cells, skeletal myoblasts and postnatal neuronal stem cells (22–24). However, much less is known about the impact of PANX1 signaling in the early embryo or in pluripotent stem cells (25).

Recent reports have revealed that PANX1 is highly expressed at the earliest stages of human development, and localizes to the plasma membrane of human oocytes as well as 2- and 4-cell stage human embryos (26). The high expression of PANX1 in human oocytes and embryos suggests a fundamental role for PANX1 in human development (25,27,28). Indeed, several human disease-causing germline *PANX1* variants have now been identified. The first human patient identified to harbor a homozygous genetic variant in *PANX1* (PANX1-R217H) suffers from a staggering number of maladies in several of the organs most highly enriched in PANX1, including severe neurological deficits and primary ovarian failure (28). This mutation was shown to decrease PANX1 channel function while not affecting trafficking. Recently, four independent families were reported in which different heterozygous *PANX1* variants cause female infertility due to primary oocyte death (26). These four human variants interfered with PANX1 posttranslational modification and plasmamembrane trafficking, decreased PANX1 protein abundance in cells, and compromised channel function.

Because PANX1 is expressed at the very earliest stages of human development, and because human mutations negatively impact human oocyte survival, we sought to uncover whether PANX1 also impacts human pluripotent stem cells or stem cell fate decisions. Here we find that PANX1 protein is expressed at the cell surface of human iPSCs. *PANX1* knockout iPSCs generated through CRISPR-Cas9 gene ablation appear morphologically indistinguishable from control. Interestingly, we find enhanced representation of endodermal and mesodermal cells from spontaneously differentiated *PANX1−/−* iPSCs exhibit compared to control. Therefore, we conclude that PANX1 protein expression influences PSC commitment to the three embryonic germ layers.

## Materials & Methods

### Induced Pluripotent Stem Cell lines

All studies were approved by the Human Ethics Research Board (HREB # 2018.210). Female wildtype human induced pluripotent stem cells (iPSCs) were created from dermal fibroblasts isolated from an apparently healthy 30-year-old female as described in Esseltine *et al*., 2017 (29) and obtained through a material transfer agreement with The University of Western Ontario. A wildtype male iPSC line (GM25256) was purchased from the Coriell Institute for Medical Research (Cat# GM25256, Coriell, Camden, NJ, USA).

iPSCs were routinely cultured as colonies in feeder-free conditions in a humidified 37°C cell culture incubator buffered with 5% CO_2_ and atmospheric oxygen. iPSCs were grown on dishes coated with Geltrex (Cat# A141330, ThermoFisher, Waltham, MA, USA) and fed daily with Essential 8 medium (Cat# A1517001, ThermoFisher). Colonies were passaged every 4-5 days when they exhibited tight cell packing, smooth borders, and phase-bright smattering at colony centers. Individual iPSCs within the colonies exhibited prominent nucleoli and high nucleus-to-cytoplasm ratio as is characteristic for human pluripotent stem cells (10,30). For passaging, iPSCs were incubated with gentle cell dissociation buffer (Cat# 13151014, ThermoFisher) at room temperature until colonies were visibly broken apart, approximately 3-5 minutes (31). Gentle cell dissociation buffer was then replaced with 1 mL of Essential 8 to stop the reaction. Colonies were then scraped from the dish surface and broken into small aggregates of cells (roughly 50 – 200 µm in diameter). The resultant aggregates were seeded into fresh Geltrex-coated wells containing Essential 8 at split ratios of 1:5 to 1:20. iPSCs were maintained in culture for 20 weeks after thawing at which point the culture was terminated and a fresh vial of low-passage iPSCs was thawed from the liquid nitrogen. We confirmed our iPSC cell banks have normal copy number at various mutation hotspots using the hPSC Genetic Analysis Kit (Cat # 07550, STEMCELL Technologies, Vancouver, BC, CAN).

Single cell iPSC passaging was achieved using StemPro Accutase (Cat# A1110501, ThermoFisher). iPSCs were treated with Accutase at 37°C for 8-10 minutes and triturated to create a single cell suspension. Single cells were plated in medium supplemented with the Rho-associated kinase inhibitor (ROCKi), Y-27632 (Cat# Y-5301, LC Laboratories, Woburn, MA, USA) to promote single cell iPSC survival (32).

### CRISPR-Cas9 gene ablation

*PANX1* knockout iPSCs were created as described previously (33). Briefly, iPSCs were transfected using the Mirus TransIT®-LT1 Transfection Reagent (Cat# MIR-2304, Mirus Bio LLC, Madison, WI, USA) with the pSpCas9(BB)-2A-GFP plasmid (Cat# 48138, Addgene, Cambridge, MA, USA) according to the manufacturer’s instructions (34). The sgRNAs were designed using the Sanger Institute CRISPR finder (http://www.sanger.ac.uk/htgt/wge/) and were selected based on their low to no exonic off-target predictions (human *PANX1*: Sanger sgRNA ID 1087081842 (5’-GCTGCGAAACGCCAGAACAG-3’)). After transfection, GFP-expressing single cells were sorted using fluorescence activated cell sorting (FACS) and re-plated at low density to permit easy isolation of individual clones. The resulting individual clones were examined for ablation of the target gene at the genomic level via PCR and Sanger sequencing while ablation of the PANX1 proteins were assessed via immunofluorescence, Western blotting, and flow cytometry.

### Embryoid Body Generation for Spontaneous Differentiation

Embryoid bodies (EBs) of 9000 cells each were created in 96-well round-bottom plates coated with 1% agarose prepared in deionized water to confer a non-adherent surface which promotes iPSC self-aggregation (35,36). A single-cell iPSC suspension was created via Accutase dissociation as described above and re-suspended in Essential 6, which lacks the essential pluripotency factors TGFβ and FGF2 (Cat# A1516401, ThermoFisher), supplemented with 10 µM Y-27632 to promote cell survival (37). Essential 6 media was replenished every other day to promote spontaneous differentiation.

### Monolayer Spontaneous Differentiation

Single cell-dissociated iPSCs were seeded onto Geltrex-coated dishes at a density of 50,000 viable cells per cm^2^. Spontaneous differentiation was initiated through Essential 6 as described above. Medium was changed every other day until Day 10 when cells were dissociated with Accutase as described above to generate single cells for flow cytometric analysis.

### Monolayer Directed Differentiation to the Three Germ Layers

Directed differentiation into the three germ layers was achieved using the STEMdiff™ Trilineage Differentiation Kit (Cat# 05230, STEMCELL Technologies) according to the manufacturer’s instructions.

### Quantitative reverse transcription polymerase chain reaction (qPCR)

Undifferentiated iPSCs, along with differentiated cells and embryoid bodies were collected for RNA extraction and qPCR gene expression analysis. RNA was extracted using the PureLink™ RNA isolation kit (Cat # 12183018A, ThermoFisher) with DNase I treatment according to the manufacturers’ instructions. Purified RNA was quantified using a NanoDrop™ 2000 spectrophotometer (Cat# ND-2000, ThermoFisher), and stored at −80°C until use. High quality RNA was identified by a ʎ260/280 of ≥ 2.0 and ʎ260/230 of ≥ 2.0.

RNA was converted into complementary DNA (cDNA) using the High-Capacity cDNA Reverse Transcription Kit (Cat# 4368814, ThermoFisher) according to the manufacturer’s instructions. Typically, 500 ng of RNA were used per cDNA reaction. The resulting cDNA was stored at −30°C until use.

Quantitative RT-PCR (qPCR) was performed using intercalating dye chemistry (38). Oligonucleotide sets were designed for specific target amplification and minimal primer dimer formation using NCBI Primer-BLAST (NIH, Bethesda, MD, USA; https://www.ncbi.nlm.nih.gov/tools/primer-blast/) and IDT’s Oligo Analyzer Tool (IDT, Newark, NJ, USA) and are listed in Table 1. Bio-Rad SsoAdvanced™ Universal SYBR® Green Supermix (Cat# 1725274, Bio-Rad, Hercules, CA, USA) was utilized and oligonucleotides (all from IDT) were used at 10 µM in each reaction. Standard run time cycling parameters were as follows: one cycle of 50°C for 2 minutes, one cycle of 95°C for 30 seconds, 40 cycles of 95°C for 10 seconds, 60°C for 1 minute, followed by a melt curve from 60°C to 95°C. Data was analyzed using QuantStudio™ real-time PCR software (Version1.3, ThermoFisher). Gene expression for each sample were normalized to the reference gene (GAPDH and/or 18S) to generate a deltaC_T_ (ΔC_T_) (39). Samples where C_T_ values were ≥ the C_T_ value of the no-template control were considered qPCR non-detects and were excluded from further analysis.

**Table 1.**
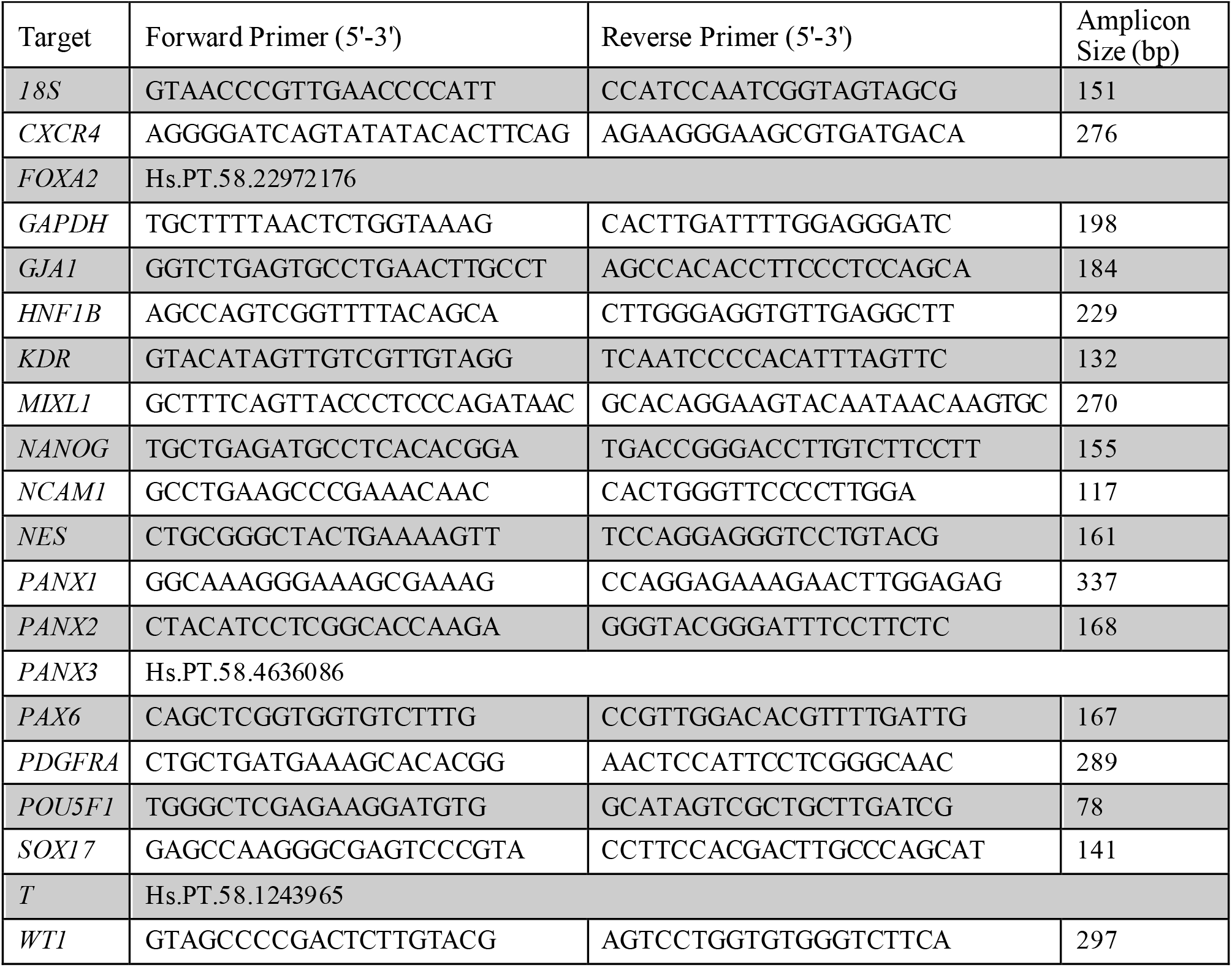
Oligo Sets for qPCR.

Heatmaps were generated from the average 2^−ΔCT^ value from each condition using R Studio (version 3.6.1) software with the ggplots.2 package and row scaling. Fold change expression of genes relative to a control sample (such as starting iPSC cell population or wildtype cells) were evaluated using the ΔΔC_T_ method as described in Schmittgen and Livak, 2008 (39). Fold change results from the average of 2-3 technical replicates were plotted in GraphPad PRISM (Version 6.07, GraphPad, San Diego, CA, USA).

### Immunofluorescence

Embryoid bodies and monolayer cultures were fixed in 10% buffered formalin (Cat# CA71007-344, VWR, Radnor, PA, USA) for 1 hour at room temperature. Fixed EBs were cryogenically prepared and immunolabelled according to the methodology described in STEMCELL Technologies’ Document #27171, Version 1.0.0, Nov 2019. In summary, EBs were first dehydrated in Ca^2+^/Mg^2+^-free PBS supplemented with 30% (w/v) sucrose for 1-4 days at 4°C until the EBs sank. Dehydrated EBs were then incubated for 1 hour at 37°C in gelatin embedding solution consisting of 10% (w/v) sucrose and 7.5% (w/v) gelatin prepared in Ca^2+^/Mg^2+^-free PBS. The EBs were then transferred to a cryopreservation mould and snap frozen in a slurry of dry ice and isopentane followed by cryosectioning at 14 µm slice thickness. For antigen retrieval, slides were placed into a plastic container with citrate buffer, pH 6.0: 0.294% tri-sodium citrate (dihydrate) (Cat# A12274, Alfa Aesar, Tewksbury, MA, USA) with 0.05% Tween®20. Samples were heated in a rice steamer for 20 minutes. Immunostaining with primary antibodies, dyes, and stains indicated in Table 2 was performed as described in the figure legends. AlexaFluor-conjugated secondary antibodies were all purchased from ThermoFisher. Slides were mounted using Mowial®488 reagent with 1,4-diazabicyclo[2.2.2]octane (DABCO) antifade compound (40).

**Table 2.**
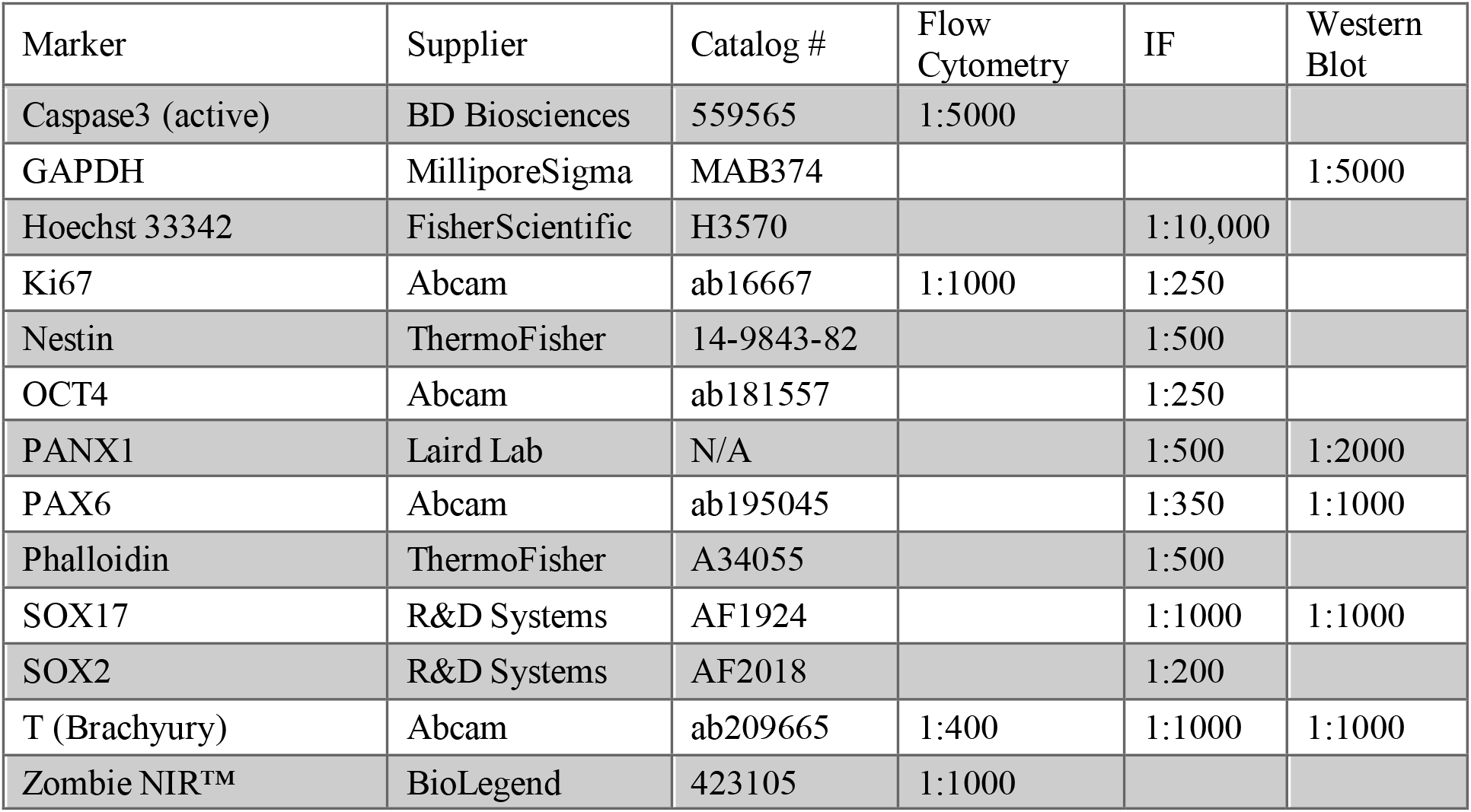
Primary Antibodies and Dyes.

| Marker | Supplier | Catalog # | Flow Cytometry | IF | Western Blot |
| --- | --- | --- | --- | --- | --- |
| Caspase3 (active) | BD Biosciences | 559565 | 1:5000 |  |  |
| GAPDH | MilliporeSigma | MAB374 |  |  | 1:5000 |
| Hoechst 33342 | FisherScientific | H3570 |  | 1:10,000 |  |
| Ki67 | Abcam | ab16667 | 1:1000 | 1:250 |  |
| Nestin | ThermoFisher | 14-9843-82 |  | 1:500 |  |
| OCT4 | Abcam | ab181557 |  | 1:250 |  |
| PANX1 | Laird Lab | N/A |  | 1:500 | 1:2000 |
| PAX6 | Abcam | ab195045 |  | 1:350 | 1:1000 |
| Phalloidin | ThermoFisher | A34055 |  | 1:500 |  |
| SOX17 | R&D Systems | AF1924 |  | 1:1000 | 1:1000 |
| SOX2 | R&D Systems | AF2018 |  | 1:200 |  |
| T (Brachyury) | Abcam | ab209665 | 1:400 | 1:1000 | 1:1000 |
| Zombie NIR™ | BioLegend | 423105 | 1:1000 |  |  |

### Phase contrast imaging

Phase contrast images of monolayer cells and embryoid bodies were taken on a Zeiss AxioObserver microscope using 5X/0.12 NA A-Plan and 10X/0.25 NA Ph1 objectives. Images from these microscopes were taken in 8-bit greyscale using an Axiocam MRm camera and AxioVision Version 4.8.2 software. All phase contrast imaging equipment is from Carl Zeiss Microscopy (Jena, DEU).

### Confocal microscopy

Fluorescent confocal images were acquired on either an Olympus Fluoview FV10i—W3 confocal microscope (Olympus, Tokyo, JPN) fitted with a 10X/0.4, or 60X/1.2 NA lens and Fluoview version 2.1.17 software. The following lasers were used to visualize fluorophores: DAPI/Hoechst (405 nm laser); Alexa Fluor 488 (473 nm laser); Alexa Fluor 555 (559 nm laser); Alexa Fluor 647 (635 nm laser). Other images were acquired using an Olympus Fluoview FV1000 confocal microscope fitted with 10X/0.4 NA, 20X/0.75NA or 40X/0.95NA and the following lasers: 405 nm, 458 nm, 568 nm, 633 nm. Images were analyzed using FIJI open source software (41). Fluorescent confocal images were occasionally subjected to brightness/contrast enhancement.

### Flow Cytometry

Flow cytometry was performed on a CytoFLEX (Beckman Coulter, Brea, CA, USA) flow cytometer. Antibodies for flow cytometry were titrated over a range of concentrations prior to use. The following controls were included in all flow cytometry runs: unstained control, fluorescence-minus-one (FMO) controls, secondary-only controls, and single-color compensation controls for fluorochromes. UltraComp compensation beads (Cat# 01-2222-43, ThermoFisher) were used with antibodies raised in mice.

Live single-cell suspensions were labelled with Zombie NIR™ fixable viability dye (Cat# 423105, BioLegend®, San Diego, CA, USA) to eliminate dead cells during the analysis stage. Next, the cells were fixed in 10% buffered formalin for 10 minutes at 4°C in the dark. After fixation, the cells were permeabilized (Ca^2+^Mg^2+^-free PBS with 0.5% w/v BSA supplemented with 0.1% Triton X-100) for 15 minutes at room temperature in the dark. Primary and secondary antibodies (used at dilutions according to Table 2) were incubated for 30 minutes at 4°C in the dark. Flow cytometric analysis was performed using FlowJo software (version 10.7.1).

### SDS-PAGE & Western Blot

Cells were lysed with a solution comprising 50mM Tris-HCl pH 8, 150 mM NaCl, 0.02% NaN_3_, 1% Triton X-100, 1 mM NaVO_4_, 10 mM NaF, 2 µg/mL leupeptin, and 2 µg/mL aprotinin. Soluble proteins were separated using SDS-PAGE and transferred to a 0.45 µm nitrocellulose membrane (Cat# 1620115, Bio-Rad). Primary antibodies (Table 2) were prepared in tris buffered saline + Tween20®(TBST) + 3% skim milk and incubated overnight at 4°C. Secondary antibodies conjugated to HRP were prepared in TBST + 3% skim milk and incubated for 1 hour at room temperature. Proteins were visualized with Bio-Rad Clarity™ Western ECL Substrate (Cat# 1705061, Bio-Rad) using a GE ImageQuant LAS 400 (Cat# 28955810, GE Healthcare, Chicago, IL, USA).

### Statistics

Statistical analysis was performed in GraphPad PRISM Version 6.07. Error bars depict ± standard error of the mean (SEM) when n ≥ 3 biological replicates (independent experiments) unless otherwise stated. Statistical significance for comparisons between 2 groups was determined by unpaired Student’s *t*-test. Statistical significance for comparisons between 3 or more groups was determined by Analysis of Variance (ANOVA) followed by a Tukey’s multiple comparisons test. * *p* < 0.05, ** *p* <0.01, *** *p* < 0.001.

## Results

### Human iPSCs express pannexin 1

Previous research has indicated that human pluripotent stem cells express transcripts of all three pannexin family members (25). Our qPCR analysis suggests that human iPSCs express mRNA for *PANX1* and *PANX2*, however we are unable to detect *PANX3* transcripts in these cells (Figure 1A). qPCR also reveals that *PANX1* gene expression is significantly higher in iPSCs compared to the dermal fibroblasts from which they were derived (Figure 1B). Flow cytometry confirms that iPSCs express PANX1 protein (Figure 1C) while immunofluorescence localizes PANX1 primarily to the cell surface of human iPSCs with lesser populations of intracellular staining (Figure 1D). Although we identified *PANX2* mRNA in our iPSCs, we were unable to find a reliable antibody to detect the PANX2 protein. Therefore, we focused our subsequent studies on PANX1.

**Figure 1.**
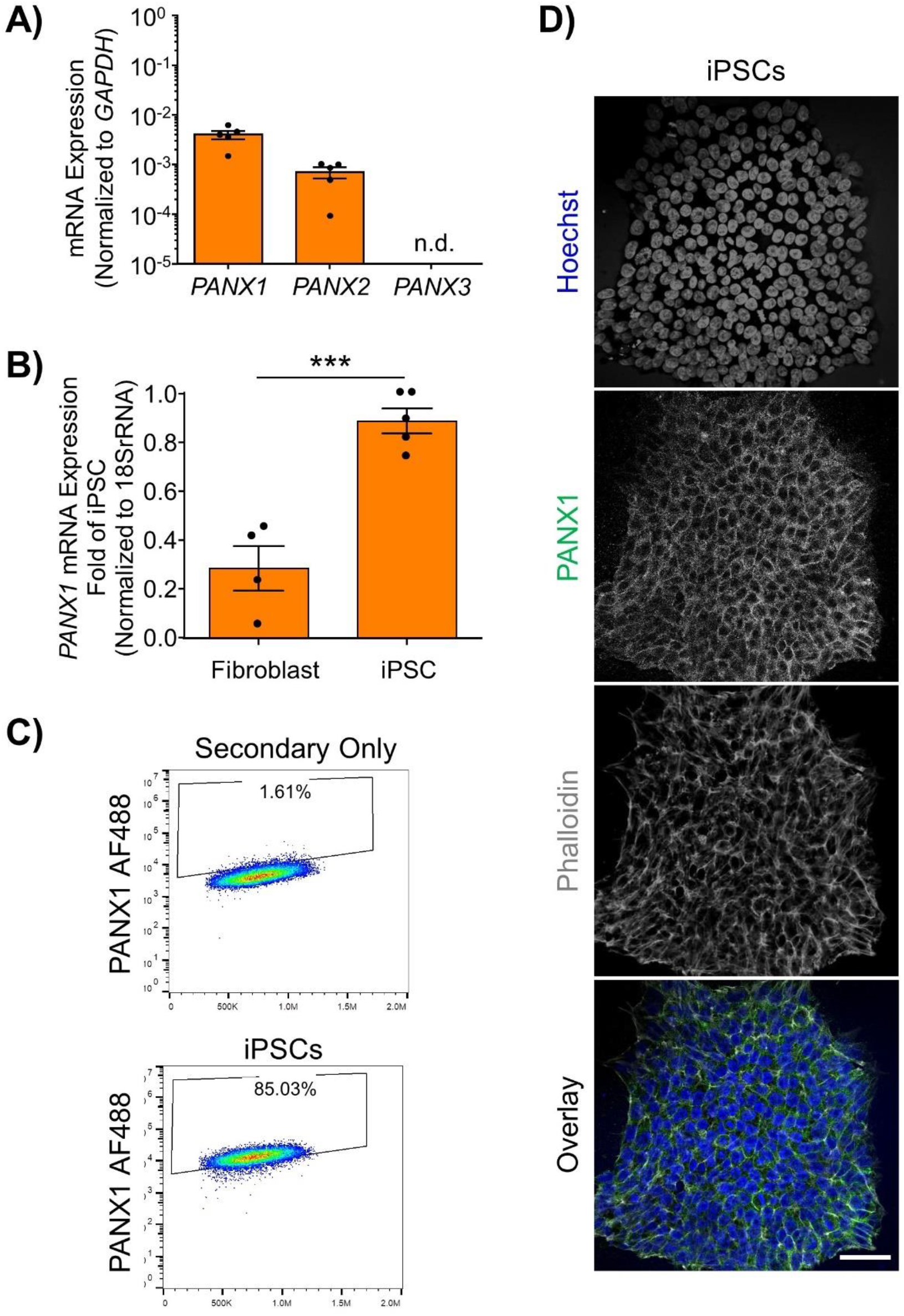
Human iPSCs express pannexin 1. **(A)** Quantitative RT-PCR (qPCR) demonstrates the presence of *PANX1* and *PANX2* transcripts in human iPSCs, but not *PANX3*. n.d. qPCR non-detect. **(B)** PANX1 mRNA is significantly upregulated in human iPSCs compared to the dermal fibroblasts from which they were derived. Data represent the standard error of the mean of 5 independent experiments. ***, p < 0.001 as assessed through Student’s T-test. **(C, D)** Pannexin 1 protein expression in human iPSCs is demonstrated through **(C)** flow cytometry and **(D)** immunofluorescence. PANX1 (green); Nuclei (Hoechst, blue); Actin (Phalloidin, grey). Scale bar = 50 μm.

### PANX1−/− iPSCs are morphologically comparable to WT

Because *PANX1* mRNA is upregulated after iPSC reprogramming, and human *PANX1* mutations are linked to primary human oocyte death, we sought to determine whether PANX1 was essential for human iPSC survival, growth, or pluripotency. CRISPR-Cas9 was used to genetically ablate *PANX1* in iPSCs (Figure 2). The resulting clonal knockout iPSCs have a single base pair deletion in the third *PANX1* exon resulting in a frameshift mutation and up to 15 early stop codons within the *PANX1* transcript (Figure 2A). At the protein level, the mutation alters the amino acid sequence starting from the second transmembrane domain (Figure 2A). Western blot analysis shows PANX1 protein in control iPSCs expressed as three distinct bands relating to the three glycosylation states where Gly2 corresponds with complex carbohydrate modification, Gly1 is the addition of a high mannose species, and Gly0 PANX1 lacks glycosylation (Figure 2B) (42). After CRISPR-Cas9 gene ablation, *PANX1−/−* iPSCs no longer express PANX1 protein as shown through Western blot, flow cytometry, and immunofluorescence (Figure 2B-D). *PANX1−/−* iPSCs appear morphologically indistinguishable from control cells and continue to grow as large colonies of tightly packed cells characteristic of human pluripotent stem cells (Figure 3A). *PANX1−/−* iPSCs continue to express similar amounts of the pluripotency markers *POU5F1* (encoding for OCT4) and *NANOG* compared to control iPSCs (Figure 3B). Furthermore, *PANX1−/−* iPSCs do not upregulate *PANX2, PANX3* nor *GJA1* (encoding for Cx43) in response to loss of the PANX1 protein (Figure 3C). Flow cytometry demonstrates equivalent expression of the proliferation marker, Ki67 as well as equally low expression of the apoptosis marker, cleaved caspase 3 (CC3) (Figure 3D). Therefore, *PANX1* genetic ablation does not appear to negatively impact human iPSC survival, proliferation, morphology, or pluripotency marker expression.

**Figure 2.**
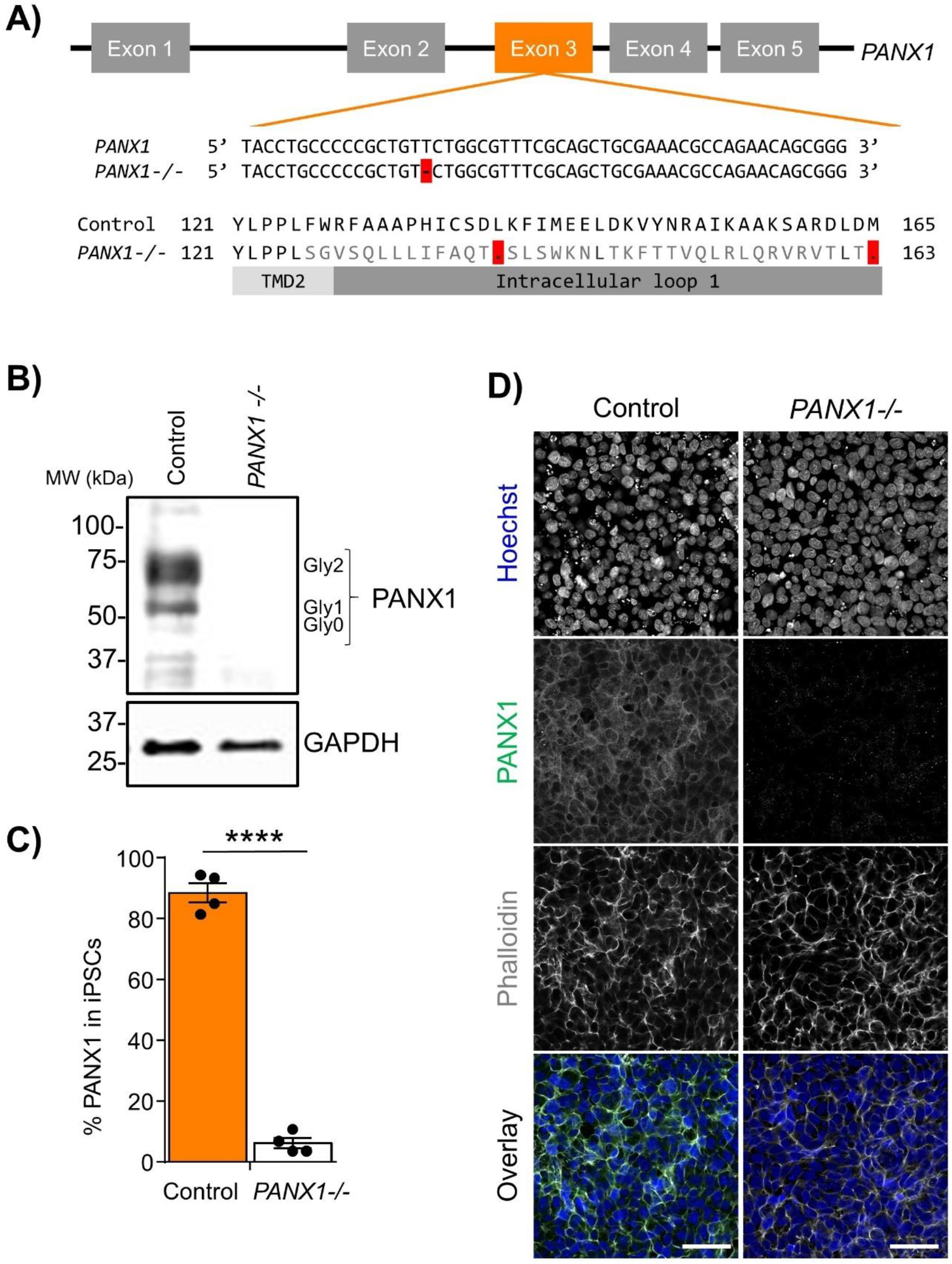
*PANX1* CRISPR-Cas9 gene ablation in human iPSCs. **(A)** CRISPR-Cas9 guide RNA was designed targeting the third exon of human *PANX1* gene. Resulting CRISPR-Cas9 engineering resulted in a single base pair deletion, thus disrupting the reading frame. **(B)** Control human iPSCs express PANX1 protein which runs on a Western blot as several discreet band sizes corresponding with the non-glycosylated protein (Gly0), high mannose (Gly1) and complex carbohydrate addition (Gly2). After CRISPR-Cas9 gene ablation, human iPSCs no longer express PANX1 protein as shown through **(B)** Western blot, **(C)** flow cytometry and **(D)** immunofluorescence. PANX1 (green); Nuclei (Hoechst, blue); Actin (Phalloidin, grey). Data represent the standard error of the mean of 4 independent experiments. ****, p < 0.0001. Scale bar = 50 μm.

**Figure 3.**
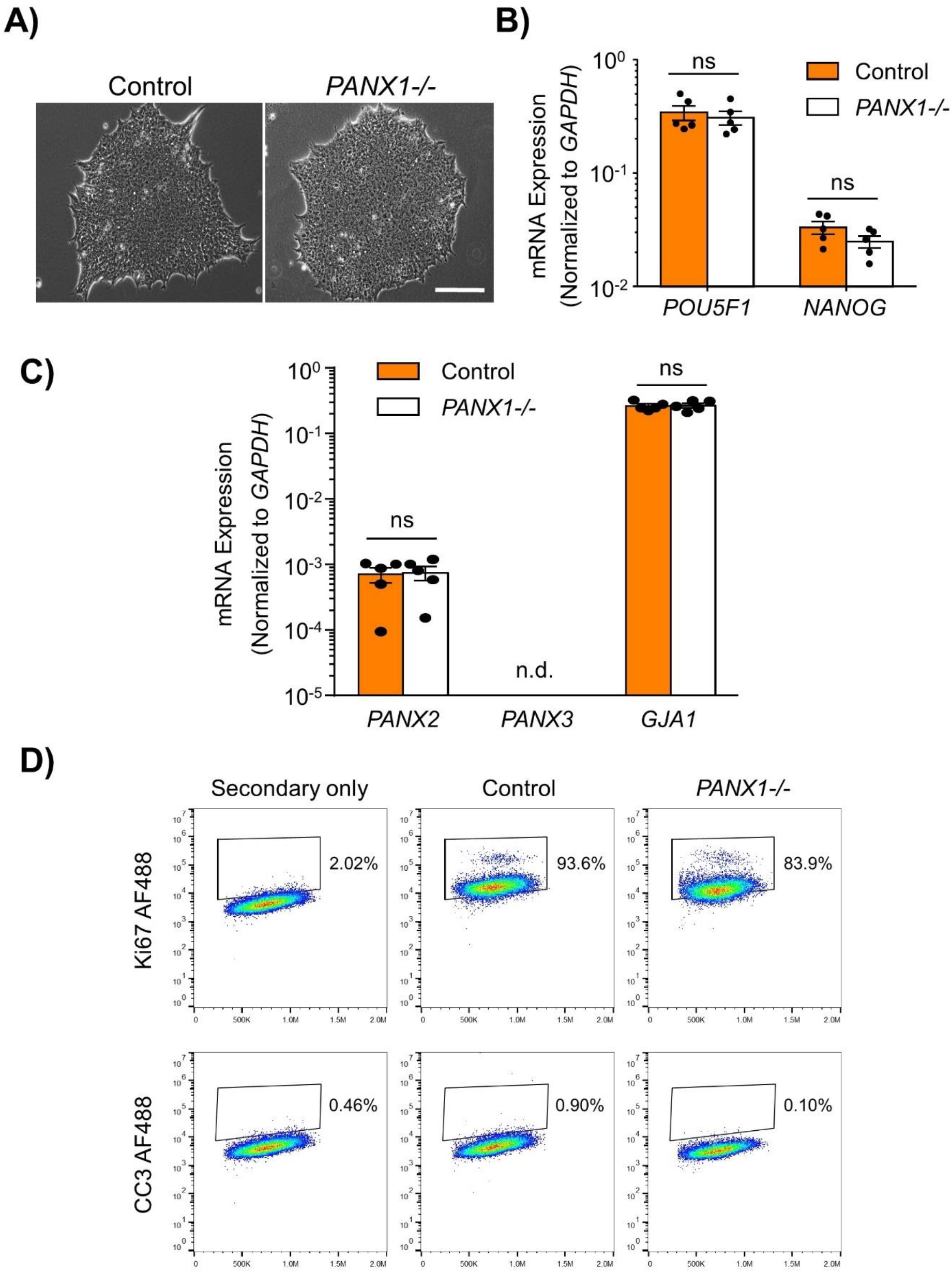
*PANX1−/−* iPSCs are morphologically comparable to control. **(A)** Phase contrast micrographs demonstrate that control and *PANX1−/−* iPSCs each grow as large colonies of tightly packed cells with refractive borders and little differentiation. Scale bar = 200 μm. qPCR analysis of **(B)** the key pluripotency genes *POU5F1* (encoding for OCT4) and *NANOG* as well as **(C)** *PANX2, PANX3* or *GJA1* (encoding for Cx43) expression in control and *PANX1−/−* iPSCs. Data represent the standard error of the mean of 5 independent experiments. ns, non-significant. n.d. qPCR non-detect. **(D)** Flow cytometry evaluation of the proliferation marker Ki67 as well as the apoptotic marker cleaved caspase 3 (CC3) in control and *PANX1−/−* iPSCs.

### Pannexin 1 is alternatively glycosylated and differentially localized in cells from the three embryonic germ layers

Although PANX1 genetic ablation was well tolerated in human iPSCs, we investigated whether this protein plays a role in cell fate specification. Given the wide expression of pannexin 1 across many tissues of the body, we examined PANX1 expression in each of the three germ layers: endoderm, ectoderm, and mesoderm (Figure 4). PANX1 was indeed expressed in each germ layer as shown through qPCR and Western blot (Figure 4A, B). Similar to what was observed in undifferentiated iPSCs, densitometric analysis of the three glycosylation species revealed that 37.15 ± 2.47% of ectoderm PANX1 is fully glycosylated while 25.59 ± 2.91% exists as high mannose and 37.24 ± 3.28% is unglycosylated (Figure 4C). Interestingly, PANX1 in mesoderm cells is significantly more glycosylated than iPSCs (61.25 ± 1.29% Gly2, 22.71 ± 7.58% Gly1 and 16.01 ± 6.43% Gly0). On the other hand, endoderm cells appeared to have a significant reduction in the Gly1 and Gly2 species. Endodermal PANX1 exists as 61.09 ± 5.84% unglycosylated species while only 22.38 ± 2.35% is fully glycosylated with complex carbohydrate species (Figure 4C). Because glycosylation has been reported to play a role in PANX1 trafficking to the plasma membrane, we also evaluated the subcellular distribution of PANX1 in the three germ lineages. Immunofluorescence localizes PANX1 protein to the cell surface in Nestin positive ectoderm cells and Brachyury positive mesoderm cells (Figure 5, inset). However, PANX1 was primarily localized to intracellular compartments of SOX17 positive endoderm cells (Figure 5, inset, yellow arrowheads).

**Figure 4.**
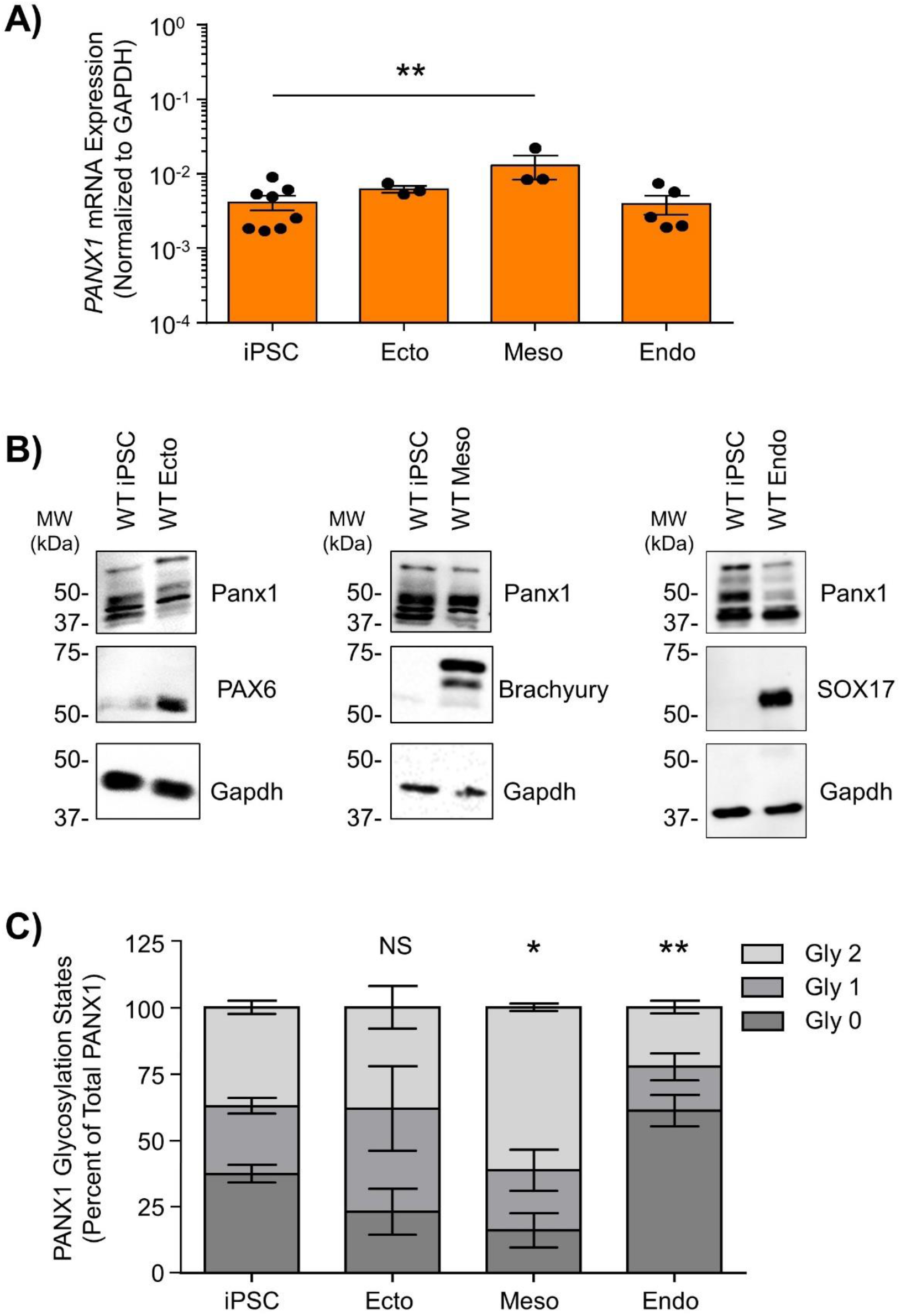
Pannexin 1 is expressed in cells from all three embryonic germ layers. **(A)** qPCR analysis of *PANX1* mRNA expression in control human iPSCs as well as after directed differentiation into each of the three embryonic germ layers, ectoderm (ecto), mesoderm (meso), and endoderm (endo). **(B** Representative Western blots and **(C)** densitometric analysis of the three glycosylation states of PANX1 in iPSCs, ectoderm, mesoderm and endoderm cells. Gly0, non-glycosylated protein; Gly1, high mannose; Gly2, complex carbohydrate addition. Data represent the standard error of the mean of 3-10 independent experiments. NS, no significance; *, p < 0.05; **, p < 0.01 compared to iPSCs.

**Figure 5.**
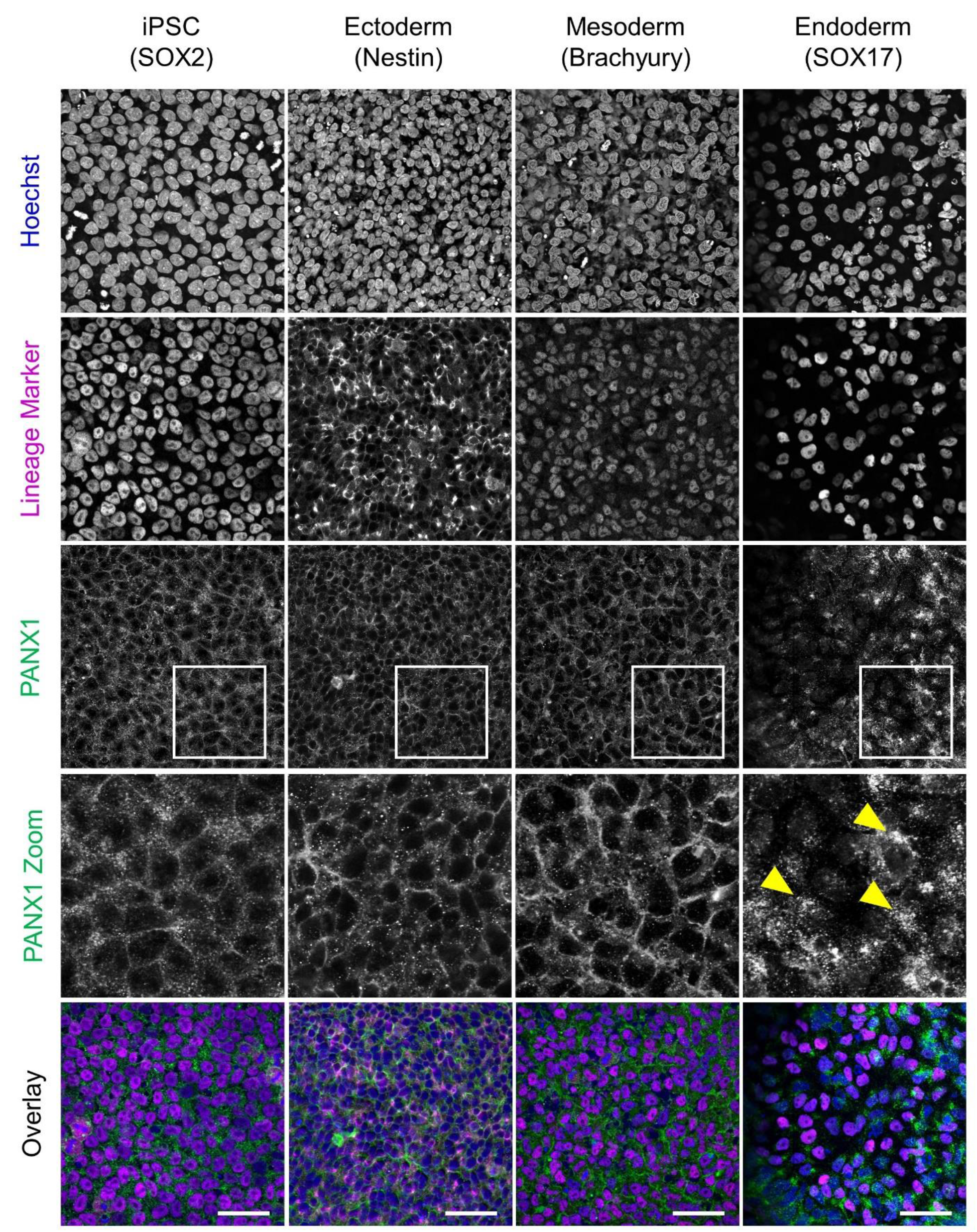
PANX1 is differentially localized in the three embryonic germ layers. Immunofluorescence evaluation of the subcellular localization of PANX1 protein (green) along with nuclei (Hoechst, blue) and lineage-specific markers (SOX2, PAX6, Brachyury, or SOX17, magenta) in control human iPSCs as well as after directed differentiation into each of the three embryonic germ layers. Inset, PANX1 is localized primarily to the cell surface in iPSCs, ectoderm and mesoderm with lesser intracellular pools. Yellow arrowheads, PANX1 is localized intracellularly in endoderm cells. Scale bar = 50 μm.

### Pannexin 1 knockout embryoid bodies exhibit skewed lineage specification

Spontaneous differentiation enables passive, cell-guided cell fate specification and generally results in cell populations from all three germ layers (ectoderm, endoderm, and mesoderm). In order to determine whether loss of PANX1 altered inherent lineage specification of human iPSCs, we investigated the spontaneous differentiation potential of control and *PANX1−/−* iPSCs cultured either as monolayers or as 3-dimensional embryoid bodies (EBs) (Figure 6A). Control and *PANX1−/−* iPSCs self-aggregated into embryoid bodies of comparable size and shape after 24 hours in culture (Figure 6B). As expected, both control and *PANX1−/−* EBs downregulated genes associated with undifferentiated state (*POU5F1* and *NANOG*) relative to starting iPSCs, indicating that the cells within the EBs were losing pluripotent stemness and were committing to downstream lineages (Figure 6C). Comprehensive gene expression analysis shows altered expression of genes associated with each of the 3 germ layers in *PANX1−/−* EBs compared to control (Figure 6C). Indeed, after 5 days of spontaneous differentiation, *PANX1−/−* EBs exhibit lower expression of genes associated with the ectoderm (*PAX6, NESTIN*) compared to control EBs at the same time points. On the other hand, expression of mesendoderm (*MIXL1*), mesoderm (*T, PDGFRA, NCAM1*) and endoderm (*SOX17, HNF1B*) markers are higher in *PANX1−/−* EBs compared to control (Figure 6C). Time course analysis suggests that at 5 days of spontaneous differentiation, expression of these endoderm and mesoderm lineage genes are higher in *PANX1−/−* EBs compared to control, and subsequently fall back to baseline expression by 14 days (Figure 7A), consistent with mesendoderm commitment and subsequent downstream lineage progression.

**Figure 6.**
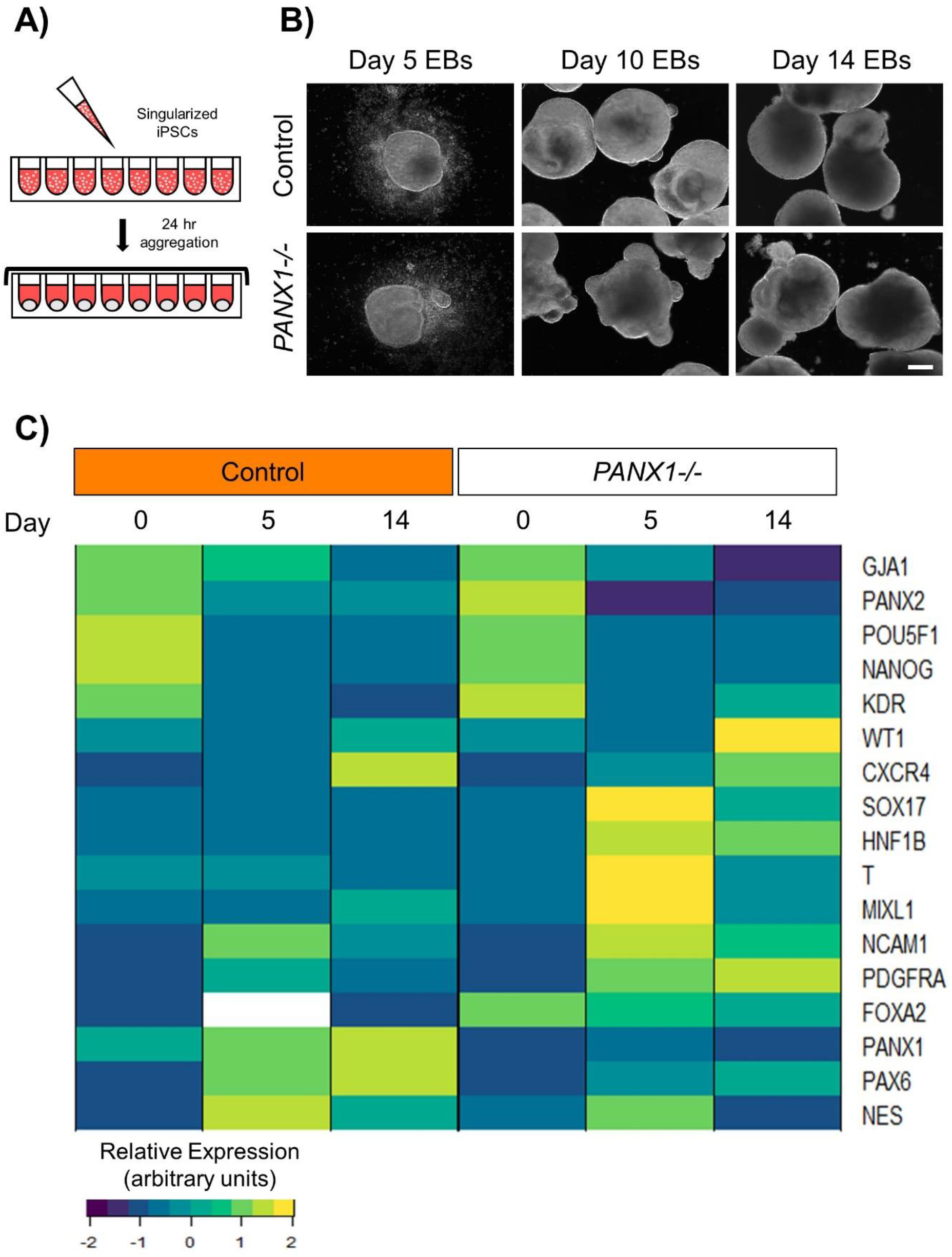
Pannexin 1 knockout embryoid bodies exhibit skewed lineage specification. **(A)** Schematic depicting embryoid body (EB) formation. **(B)** Control and *PANX1−/−* iPSCs self-aggregate and form embryoid bodies. Scale bar = 200 μm. **(C)** Gene expression analysis of control and *PANX1−/−* iPSCs (day 0) as well as embryoid bodies after 5 and 14 days of spontaneous differentiation. Heatmaps were generated from the average 2^ΔCT^ value from each condition using R Studio software with the ggplots.2 package and row clustering. Data represents the mean of 5 independent experiments at day 0 and 5, and 3 independent experiments at day 14.

**Figure 7.**
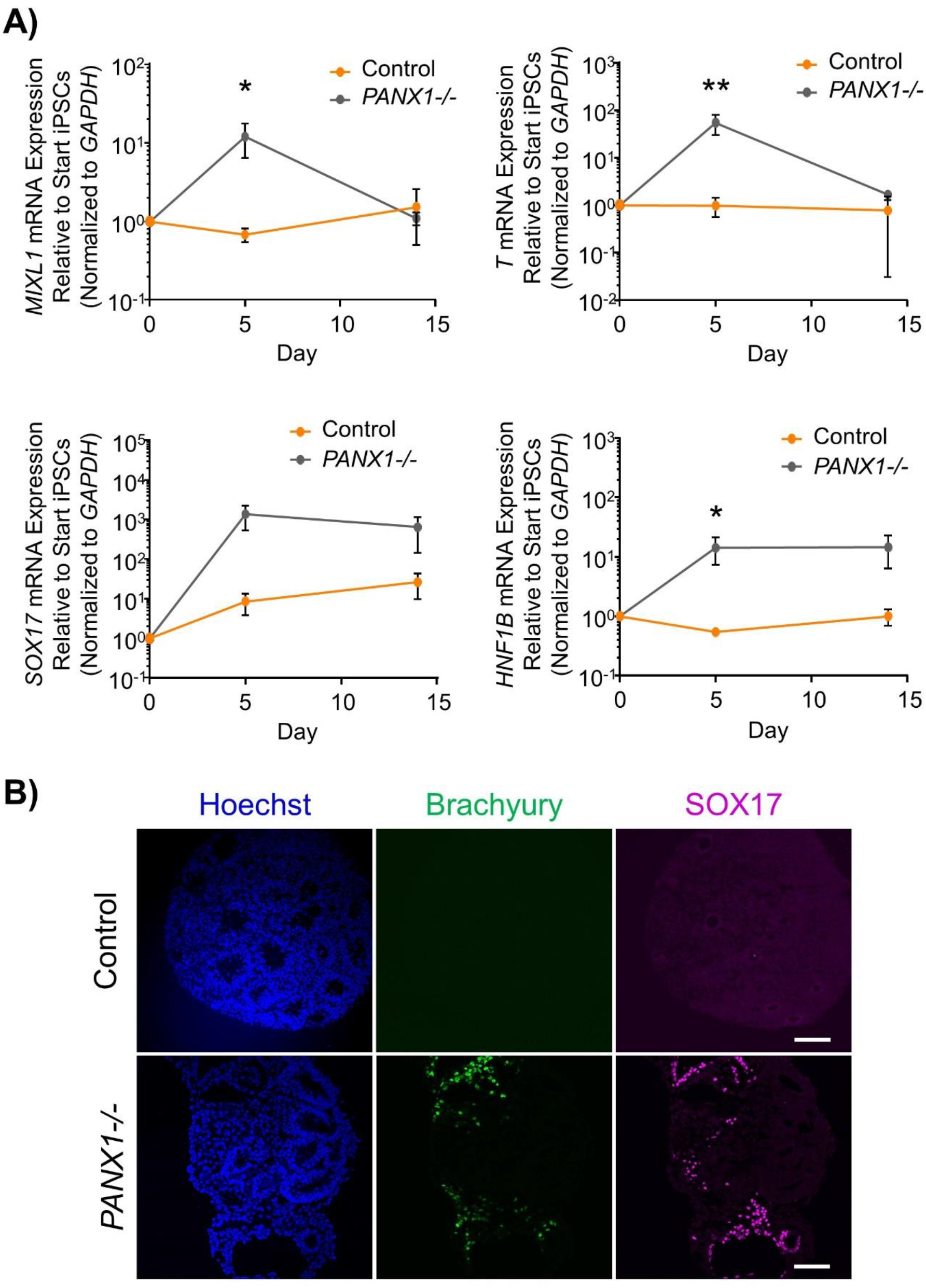
Pannexin 1 gene ablation results in skewed lineage specification of human iPSCs. **(A)** qPCR gene expression analysis of control and *PANX1−/−* iPSCs (day 0) as well as embryoid bodies after 5 and 14 days of spontaneous differentiation. *, p < 0.05; **, p <0.001 compared to control at the same timepoint. **(B)** Immunofluorescence of day 14 control and *PANX1−/−* EBs. Nuclei (Hoechst, blue); mesoderm (Brachyury, green); and endoderm (SOX17, magenta). Scale bar = 100 μm.

We corroborated our qPCR data using immunofluorescence imaging of control and *PANX1−/−* embryoid bodies (Figure 7B). Consistent with our gene expression analysis, we observed a greater proportion of *PANX1−/−* cells expressing Brachyury (mesoderm) and SOX17 (endoderm) relative to control (Figure 7B). Taken together, spontaneously differentiated *PANX1−/−* EBs appear to favor formation of mesoderm and endoderm germ layers with reduced capacity to for ectodermal cells.

### Exogenous pressures override PANX1−/− lineage bias

We determined above that *PANX1−/−* iPSCs exhibit apparent lineage specification bias when permitted to spontaneously differentiate. However, we also wanted to determine whether loss of PANX1 impacted directed lineage differentiation promoted through the application of exogenous growth factors and small molecules. To that end, we used the STEMdiff™ Trilineage Differentiation Kit (STEMCELL Technologies) to evaluate the ability of *PANX1−/−* iPSCs to differentiate into cells from all three germ layers in response to external pressures. Despite the demonstrated lineage biases outlined above in the passive cultures, both qPCR and Western blot demonstrate that *PANX1−/−* iPSCs effectively differentiated into cells of all three germ lineages when cultured with the Trilineage Differentiation Kit (Figure 8). Similar to what we observed in the undifferentiated iPSCs, both *PANX2* and *GJA1* (Cx43) transcripts are expressed in the three germ layers, but are not upregulated in compensation for the loss of PANX1 during directed differentiation (Figure 9). Furthermore, *PANX3* transcripts remained undetectable by qPCR in cells from all three germ layers (data not shown). Therefore, although we find that pannexin 1 influences germ layer specification, it is not essential to this process.

**Figure 8.**
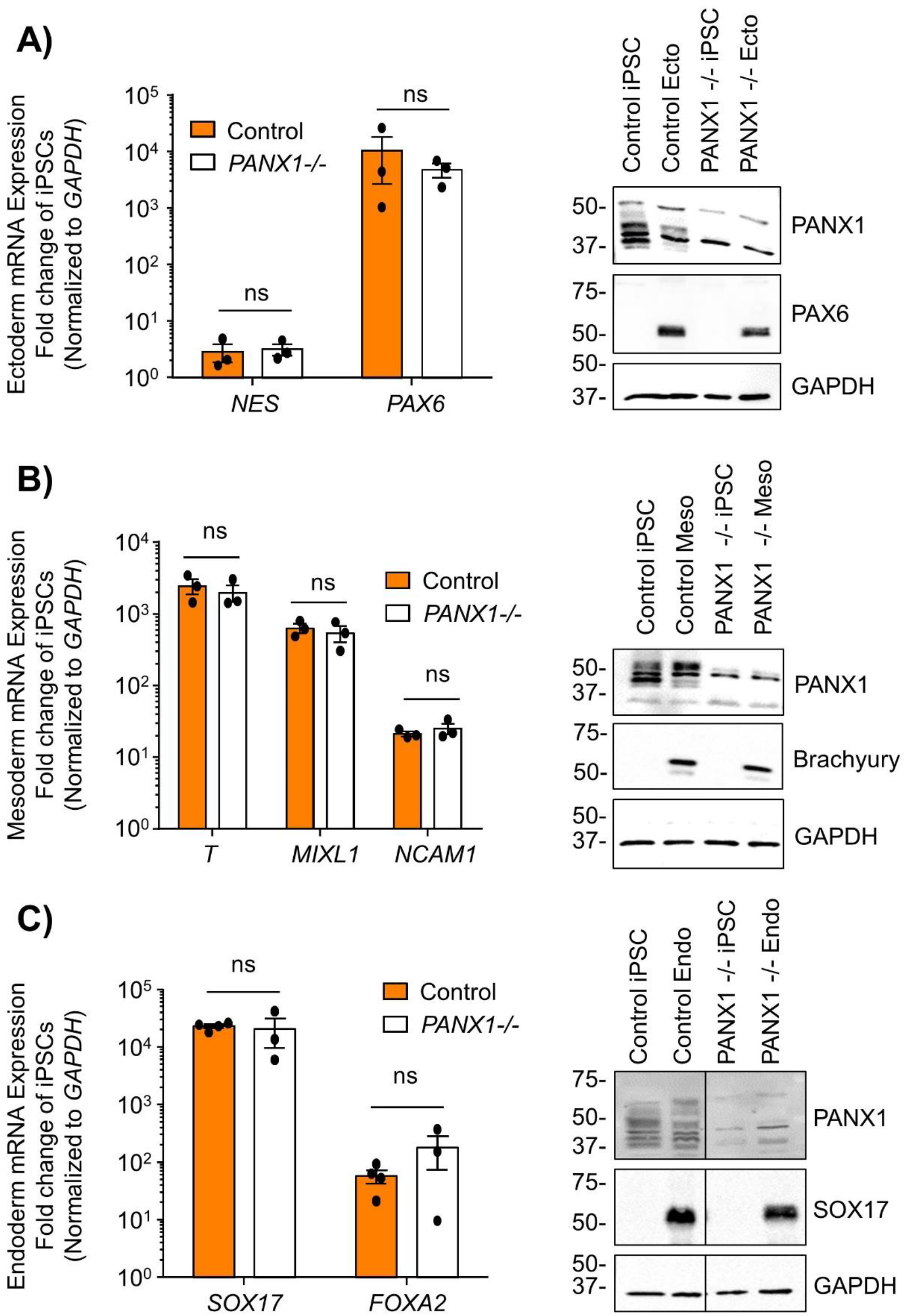
Exogenous pressures override *PANX1−/−* lineage bias. qPCR gene expression analysis and Western blot of lineage-specific markers after directed differentiation of control and *PANX1−/−* iPSCs into **(A)** Ectoderm (*NES, PAX6*), **(B)** Mesoderm (*T* (Brachyury), *MIXL1, NCAM1*) and **(C)** Endoderm (*SOX17, FOXA2*). Data represent the standard error of the mean of 3-4 independent experiments. ns, non-significant compared to control.

**Figure 9.**
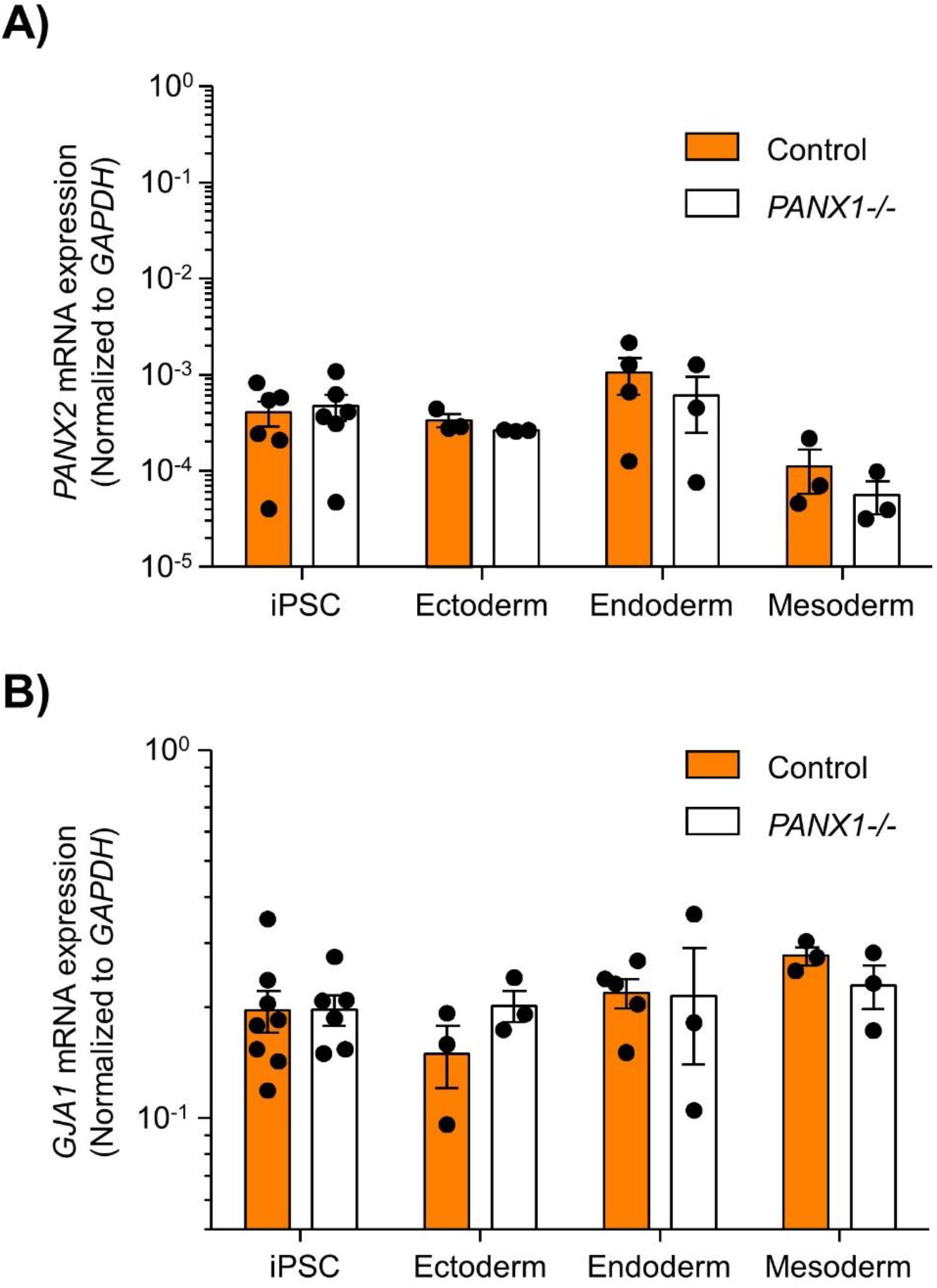
PANX2 and Cx43 do not compensate for PANX1 loss during germ lineage differentiation. qPCR analysis of **(A)** *PANX2* and **(B)** *GJA1* (Cx43) mRNA expression in directed ectoderm, mesoderm, and endoderm cultures. Data represent the standard error of the mean of 3-8 independent experiments.

## Discussion

Here we find that PANX1 protein is expressed in human iPSCs as well as in cells of all three embryonic germ lineages: ectoderm, endoderm, and mesoderm, further underlying the potential for this protein to participate in cell fate specification. Despite being dispensable for stem cell morphology, proliferation, and survival, PANX1 does contribute to cell fate specification with lower ectodermal representation and exaggerated mesendoderm cell abundance in spontaneously differentiated *PANX1−/−* cultures compared to control.

The human protein atlas reports that pannexin 1 is widely expressed in tissues arising from all three germ layers (https://www.proteinatlas.org/ENSG00000110218-PANX1/tissue) and the many mouse studies conducted around the world have highlighted how important murine Panx1 is in postnatal health and disease. For example, Panx1 is reported to exacerbate the spread of ischemic injury in mice following a stroke via the disruption of electrochemical gradients in neurons and glial cell types (43,44). Human PANX1 also has reported involvement in pathogen-mediated activation of the caspase cascade by releasing ATP which attracts phagocytic cells, resulting in the clearance of the damaged/infected cells (17,43). On the other hand, HIV can use the PANX1 channel to enter lymphocytes (45) and once inside the cell, the virus can elicit PANX1-mediated ATP release to destabilize the cell membrane and ultimately facilitate viral spread (44,46). Surprisingly, few studies have focused on human pannexin proteins or the role of pannexins in early development. We are now able to uncover how PANX1 signaling influences the earliest developmental decisions through spontaneous and directed differentiation of human iPSCs, and by modelling human tissue development through the use of PSC-derived organoids.

PANX1 is expressed in the human oocyte as well as the 2- and 4-cell stage embryo (26). Human embryonic stem cells and induced pluripotent stem cells have previously been shown to express all three pannexin transcripts (25). However, we only detect expression of *PANX1* and *PANX2* and were unable to detect *PANX3* in any of the stem cell types that we evaluated. We also show that human iPSCs express pannexin 1 protein, which is concentrated at the cell surface of undifferentiated iPSCs with lesser amounts of intracellular PANX1 localization. Furthermore, PANX1 expression and localization persist in cells of all three embryonic germ layers. Interestingly, while PANX1 was primarily at the cell surface of iPSCs, ectoderm and mesoderm cells, it was intracellularly localized in endoderm cells. This intracellular localization was accompanied by a significant reduction in the complex carbohydrate posttranslational modification as identified through Western blot. It remains to be seen what role this intracellular PANX1 pool plays in endodermal tissues, but given that our PANX1 −/− iPSCs exhibit enhanced endodermal differentiation, removing this protein is clearly beneficial for endodermal differentiation.

Most of what is currently understood about pannexins in stem cell fate specification arises from somatic (adult or tissue-resident) stem cell studies. Pannexin signaling can influence the self-renewal and differentiation of multiple somatic stem cell types including osteoprogenitor cells, skeletal myoblasts and postnatal neuronal stem cells (22–24). Panx1 and Panx2 are involved in postnatal murine neural progenitor cell maintenance and proliferation (22). Panx1 participates in murine NPC (neural progenitor cell) self-renewal while Panx2 has been implicated in postnatal murine neuronal differentiation (47,48). Meanwhile, Panx1 and Panx3 are both expressed in mesodermal tissues such as boneand cartilage and they haveboth been implicated in the regulation and commitment of resident progenitor cell populations in mice (24,49–51). Panx1 is expressed in murine bone marrow-derived stromal cells while Panx3 inhibits osteoprogenitor cell proliferation and contributes to chondrocyte differentiation (49,50). In contrast to these studies on adult progenitor cell populations, very little is understood about pannexins in pluripotent stem cells, including mouse and human ESCs or iPSCs.

Human germline *PANX1* mutations which cause decreased protein abundance and trafficking defects lead to human oocyte death and female infertility (26). This effect can be mimicked using isolated mouse oocytes injected with *PANX1* mutant complementary RNA (cRNA). On the other hand, *Panx1* knockout mice remain fertile and continue to birth average litter sizes. These observed differences in fecundity may be due to inherent species differences, or *in vitro* versus *in vivo* manipulations. We find that CRISPR-Cas9 genetic ablation of *PANX1* does not negatively impact human iPSC proliferation, survival, or morphology. It is possible that *PANX1* missense mutations are more impactful than complete ablation due to as-yet unknown gain of function properties or changes in protein partner interactions. It would be interesting to examine whether human iPSCs are amenable to insertion of human missense mutations via gene editing or whether *PANX1−/−* iPSCs can differentiate to primordial germ cells.

The most well-defined role of pannexin 1 is in the regulated release of ATP through several mechanisms including mechanical stress, membrane depolarization, changes in intracellular ion concentration and others (16). Autocrine and paracrine signaling mechanisms triggered through cellular release of ATP and ADP have reported trophic, differentiating, and immunomodulatory effects and ATP signaling has been linked to proliferation of mouse embryonic stem cell and several postnatal progenitor cell populations (13,52). Activated pannexin channels appear to play a supporting role in augmenting purinergic receptor activity through the release of extracellular nucleotides and nucleosides. Additionally, pannexin 1 has been widely implicated in cell death signaling (17,44,53). Apoptotic induction through caspase activation leads to cleavage of the PANX1 carboxyl-terminal tail, which subsequently results in release of ATP and other small molecules into the extracellular space. Although we have found that iPSCs tolerate the loss of PANX1, follow up investigations are necessary to determine whether the lineage bias that we have presented here actually results from compromised ectodermal lineage specification or due to altered proliferation or apoptosis of certain germ layers within *PANX1* null cells.

Human developmental disorders arise in part due to flawed cell fate specification which can contribute to organ and tissue dysfunction. Until we have a comprehensive understanding of human development and cell fate decisions our capacity to treat developmental disorders remains incomplete. Given the ubiquitous expression of PANX1 in adult tissues, we expect one or more aspects of human stem cell pluripotency or early lineage specification to be affected by the loss of PANX1. Future studies will uncover protein interacting partners and specific messenger molecules involved in this process. Furthermore, it will be interesting to determine whether pannexin 1 is similarly involved in downstream specification of various terminally differentiated cells or 3-dimensional organoid development.

## Acknowledgements

We thank Dr. Dale Laird for generously providing us with the control and *PANX1−/−* iPSCs as well as the human PANX1 antibody used in this study.

## Author Contributions

J.L.E. used CRISPR/Cas9 to generate the *PANX1−/−* iPSCs. R.J.N and G.A.C performed all the experiments, analyzed the data and assembled the figures. J.L.E. oversaw the project and wrote the manuscript. All authors reviewed and edited the manuscript.

## References

1. Itskovitz-Eldor, J., Schuldiner, M., Karsenti, D., Eden, A., Yanuka, O., Amit, M., Soreq, H., and Benvenisty, N. (2000) Differentiation of human embryonic stem cells into embryoid bodies compromising the three embryonic germ layers. Mol Med 6, 88–95

2. Nagy, A., Rossant, J., Nagy, R., Abramow-Newerly, W., and Roder, J. C. (1993) Derivation of completely cell culture-derived mice from early-passage embryonic stem cells. Proc Natl Acad Sci U S A 90, 8424–8428

3. Tam, P. P., and Loebel, D. A. (2007) Gene function in mouse embryogenesis: get set for gastrulation. Nat Rev Genet 8, 368–381

4. Camacho-Aguilar, E., and Warmflash, A. (2020) Insights into mammalian morphogen dynamics from embryonic stem cell systems. Curr Top Dev Biol 137, 279–305

5. Tada, S., Era, T., Furusawa, C., Sakurai, H., Nishikawa, S., Kinoshita, M., Nakao, K., and Chiba, T. (2005) Characterization of mesendoderm: a diverging point of the definitive endoderm and mesoderm in embryonic stem cell differentiation culture. Development 132, 4363–4374

6. Tan, J. Y., Sriram, G., Rufaihah, A. J., Neoh, K. G., and Cao, T. (2013) Efficient derivation of lateral plate and paraxial mesoderm subtypes from human embryonic stem cells through GSKi-mediated differentiation. Stem Cells Dev 22, 1893–1906

7. ten Berge, D., Koole, W., Fuerer, C., Fish, M., Eroglu, E., and Nusse, R. (2008) Wnt signaling mediates self-organization and axis formation in embryoid bodies. Cell Stem Cell 3, 508–518

8. Tchieu, J., Zimmer, B., Fattahi, F., Amin, S., Zeltner, N., Chen, S., and Studer, L. (2017) A Modular Platform for Differentiation of Human PSCs into All Major Ectodermal Lineages. Cell Stem Cell 21, 399–410.e397

9. Desbaillets, I., Ziegler, U., Groscurth, P., and Gassmann, M. (2000) Embryoid bodies: an in vitro model of mouse embryogenesis. Exp Physiol 85, 645–651

10. Takahashi, K., Tanabe, K., Ohnuki, M., Narita, M., Ichisaka, T., Tomoda, K., and Yamanaka, S. (2007) Induction of pluripotent stem cells from adult human fibroblasts by defined factors. Cell 131, 861–872

11. Perrimon, N., Pitsouli, C., and Shilo, B. Z. (2012) Signaling mechanisms controlling cell fate and embryonic patterning. Cold Spring Harb Perspect Biol 4, a005975

12. Kaebisch, C., Schipper, D., Babczyk, P., and Tobiasch, E. (2015) The role of purinergic receptors in stem cell differentiation. Comput Struct Biotechnol J 13, 75–84

13. Cavaliere, F., Donno, C., and D’Ambrosi, N. (2015) Purinergic signaling: a common pathway for neural and mesenchymal stem cell maintenance and differentiation. Front Cell Neurosci 9, 211

14. Glaser, T., de Oliveira, S. L., Cheffer, A., Beco, R., Martins, P., Fornazari, M., Lameu, C., Junior, H. M., Coutinho-Silva, R., and Ulrich, H. (2014) Modulation of mouse embryonic stem cell proliferation and neural differentiation by the P2X7 receptor. PLoS One 9, e96281

15. Jiang, L. H., Hao, Y., Mousawi, F., Peng, H., and Yang, X. (2017) Expression of P2 Purinergic Receptors in Mesenchymal Stem Cells and Their Roles in Extracellular Nucleotide Regulation of Cell Functions. J Cell Physiol 232, 287–297

16. Penuela, S., Gehi, R., and Laird, D. W. (2013) The biochemistry and function of pannexin channels. Biochim Biophys Acta 1828, 15–22

17. Chekeni, F. B., Elliott, M. R., Sandilos, J. K., Walk, S. F., Kinchen, J. M., Lazarowski, E. R., Armstrong, A. J., Penuela, S., Laird, D. W., Salvesen, G. S., Isakson, B. E., Bayliss, D. A., and Ravichandran, K. S. (2010) Pannexin 1 channels mediate ‘find-me’ signal release and membrane permeability during apoptosis. Nature 467, 863–867

18. Sandilos, J. K., and Bayliss, D. A. (2012) Physiological mechanisms for the modulation of pannexin 1 channel activity. J Physiol 590, 6257–6266

19. Bao, L., Locovei, S., and Dahl, G. (2004) Pannexin membrane channels are mechanosensitive conduits for ATP. FEBS Lett 572, 65–68

20. Furlow, P. W., Zhang, S., Soong, T. D., Halberg, N., Goodarzi, H., Mangrum, C., Wu, Y. G., Elemento, O., and Tavazoie, S. F. (2015) Mechanosensitive pannexin-1 channels mediate microvascular metastatic cell survival. Nat Cell Biol 17, 943–952

21. Sanchez-Arias, J. C., Liu, M., Choi, C. S. W., Ebert, S. N., Brown, C. E., and Swayne, L. A. (2019) Pannexin 1 Regulates Network Ensembles and Dendritic Spine Development in Cortical Neurons. eNeuro 6

22. Swayne, L. A., and Bennett, S. A. (2016) Connexins and pannexins in neuronal development and adult neurogenesis. BMC Cell Biol 17 Suppl 1, 10

23. Ishikawa, M., and Yamada, Y. (2017) The Role of Pannexin 3 in Bone Biology. J Dent Res 96, 372–379

24. Pham, T. L., St-Pierre, M. E., Ravel-Chapuis, A., Parks, T. E. C., Langlois, S., Penuela, S., Jasmin, B. J., and Cowan, K. N. (2018) Expression of Pannexin 1 and Pannexin 3 during skeletal muscle development, regeneration, and Duchenne muscular dystrophy. J Cell Physiol 233, 7057–7070

25. Hainz, N., Beckmann, A., Schubert, M., Haase, A., Martin, U., Tschernig, T., and Meier, C. (2018) Human stem cells express pannexins. BMC Res Notes 11, 54

26. Sang, Q., Zhang, Z., Shi, J., Sun, X., Li, B., Yan, Z., Xue, S., Ai, A., Lyu, Q., Li, W., Zhang, J., Wu, L., Mao, X., Chen, B., Mu, J., Li, Q., Du, J., Sun, Q., Jin, L., He, L., Zhu, S., Kuang, Y., and Wang, L. (2019) A pannexin 1 channelopathy causes human oocyte death. Sci Transl Med 11

27. Esseltine, J. L., and Laird, D. W. (2016) Next-Generation Connexin and Pannexin Cell Biology. Trends Cell Biol 26, 944–955

28. Shao, Q., Lindstrom, K., Shi, R., Kelly, J., Schroeder, A., Juusola, J., Levine, K. L., Esseltine, J. L., Penuela, S., Jackson, M. F., and Laird, D. W. (2016) A Germline Variant in the PANX1 Gene Has Reduced Channel Function and Is Associated with Multisystem Dysfunction. J Biol Chem 291, 12432–12443

29. Esseltine, J. L., Shao, Q., Brooks, C., Sampson, J., Betts, D. H., Séguin, C. A., and Laird, D. W. (2017) Connexin43 Mutant Patient-Derived Induced Pluripotent Stem Cells Exhibit Altered Differentiation Potential. J Bone Miner Res 32, 1368–1385

30. Thomson, J. A., Itskovitz-Eldor, J., Shapiro, S. S., Waknitz, M. A., Swiergiel, J. J., Marshall, V. S., and Jones, J. M. (1998) Embryonic stem cell lines derived from human blastocysts. Science 282, 1145–1147

31. Beers, J., Gulbranson, D. R., George, N., Siniscalchi, L. I., Jones, J., Thomson, J. A., and Chen, G. (2012) Passaging and colony expansion of human pluripotent stem cells by enzyme-free dissociation in chemically defined culture conditions. Nat Protoc 7, 2029–2040

32. Watanabe, K., Ueno, M., Kamiya, D., Nishiyama, A., Matsumura, M., Wataya, T., Takahashi, J. B., Nishikawa, S., Muguruma, K., and Sasai, Y. (2007) A ROCK inhibitor permits survival of dissociated human embryonic stem cells. Nat Biotechnol 25, 681–686

33. Esseltine, J. L., Brooks, C. R., Edwards, N. A., Subasri, M., Sampson, J., Séguin, C., Betts, D. H., and Laird, D. W. (2020) Dynamic regulation of connexins in stem cell pluripotency. Stem Cells 38, 52–66

34. Ran, F. A., Hsu, P. D., Wright, J., Agarwala, V., Scott, D. A., and Zhang, F. (2013) Genome engineering using the CRISPR-Cas9 system. Nat Protoc 8, 2281–2308

35. Dahlmann, J., Kensah, G., Kempf, H., Skvorc, D., Gawol, A., Elliott, D. A., Dräger, G., Zweigerdt, R., Martin, U., and Gruh, I. (2013) The use of agarose microwells for scalable embryoid body formation and cardiac differentiation of human and murine pluripotent stem cells. Biomaterials 34, 2463–2471

36. Friedrich, J., Seidel, C., Ebner, R., and Kunz-Schughart, L. A. (2009) Spheroid-based drug screen: considerations and practical approach. Nat Protoc 4, 309–324

37. Lin, Y., and Chen, G. (2014) StemBook. in StemBook (Community, T. S. C. R. ed.), IOS Press. pp

38. Zipper, H., Brunner, H., Bernhagen, J., and Vitzthum, F. (2004) Investigations on DNA intercalation and surface binding by SYBR Green I, its structure determination and methodological implications. Nucleic Acids Res 32, e103

39. Schmittgen, T. D., and Livak, K. J. (2008) Analyzing real-time PCR data by the comparative C(T) method. Nat Protoc 3, 1101–1108

40. Laboratory, C. S. H. (2007) Mowial-DABCO stock solution. Cold Spring Harb. Protoc.

41. Schindelin, J., Arganda-Carreras, I., Frise, E., Kaynig, V., Longair, M., Pietzsch, T., Preibisch, S., Rueden, C., Saalfeld, S., Schmid, B., Tinevez, J. Y., White, D. J., Hartenstein, V., Eliceiri, K., Tomancak, P., and Cardona, A. (2012) Fiji: an open-source platform for biological-image analysis. Nat Methods 9, 676–682

42. Penuela, S., Bhalla, R., Gong, X. Q., Cowan, K. N., Celetti, S. J., Cowan, B. J., Bai, D., Shao, Q., and Laird, D. W. (2007) Pannexin 1 and pannexin 3 are glycoproteins that exhibit many distinct characteristics from the connexin family of gap junction proteins. J Cell Sci 120, 3772–3783

43. Bond, S. R., and Naus, C. C. (2014) The pannexins: past and present. Front Physiol 5, 58

44. Penuela, S., Harland, L., Simek, J., and Laird, D. W. (2014) Pannexin channels and their links to human disease. Biochem J 461, 371–381

45. Orellana, J. A., Velasquez, S., Williams, D. W., Sáez, J. C., Berman, J. W., and Eugenin, E. A. (2013) Pannexin1 hemichannels are critical for HIV infection of human primary CD4+ T lymphocytes. J Leukoc Biol 94, 399–407

46. Séror, C., Melki, M. T., Subra, F., Raza, S. Q., Bras, M., Saïdi, H., Nardacci, R., Voisin, L., Paoletti, A., Law, F., Martins, I., Amendola, A., Abdul-Sater, A. A., Ciccosanti, F., Delelis, O., Niedergang, F., Thierry, S., Said-Sadier, N., Lamaze, C., Métivier, D., Estaquier, J., Fimia, G. M., Falasca, L., Casetti, R., Modjtahedi, N., Kanellopoulos, J., Mouscadet, J. F., Ojcius, D. M., Piacentini, M., Gougeon, M. L., Kroemer, G., and Perfettini, J. L. (2011) Extracellular ATP acts on P2Y2 purinergic receptors to facilitate HIV-1 infection. J Exp Med 208, 1823–1834

47. Swayne, L. A., Sorbara, C. D., and Bennett, S. A. (2010) Pannexin 2 is expressed by postnatal hippocampal neural progenitors and modulates neuronal commitment. J Biol Chem 285, 24977–24986

48. Wicki-Stordeur, L. E., and Swayne, L. A. (2012) Large Pore Ion and Metabolite-Permeable Channel Regulation of Postnatal Ventricular Zone Neural Stem and Progenitor Cells: Interplay between Aquaporins, Connexins, and Pannexins? Stem Cells Int 2012, 454180

49. Bond, S. R., Lau, A., Penuela, S., Sampaio, A. V., Underhill, T. M., Laird, D. W., and Naus, C. C. (2011) Pannexin 3 is a novel target for Runx2, expressed by osteoblasts and mature growth plate chondrocytes. J Bone Miner Res 26, 2911–2922

50. Ishikawa, M., Iwamoto, T., Fukumoto, S., and Yamada, Y. (2014) Pannexin 3 inhibits proliferation of osteoprogenitor cells by regulating Wnt and p21 signaling. J Biol Chem 289, 2839–2851

51. Lee, V. R., Barr, K. J., Kelly, J. J., Johnston, D., Brown, C. F. C., Robb, K. P., Sayedyahossein, S., Huang, K., Gros, R., Flynn, L. E., and Penuela, S. (2018) Pannexin 1 regulates adipose stromal cell differentiation and fat accumulation. Sci Rep 8, 16166

52. Heo, J. S., and Han, H. J. (2006) ATP stimulates mouse embryonic stem cell proliferation via protein kinase C, phosphatidylinositol 3-kinase/Akt, and mitogen-activated protein kinase signaling pathways. Stem Cells 24, 2637–2648

53. Imamura, H., Sakamoto, S., Yoshida, T., Matsui, Y., Penuela, S., Laird, D. W., Mizukami, S., Kikuchi, K., and Kakizuka, A. (2020) Single-cell dynamics of pannexin-1-facilitated programmed ATP loss during apoptosis. Elife 9

